# Reducing INDEL calling errors in whole-genome and exome sequencing data

**DOI:** 10.1101/006148

**Authors:** Han Fang, Yiyang Wu, Giuseppe Narzisi, Jason A. O’Rawe, Laura T. Jimenez Barrón, Julie Rosenbaum, Michael Ronemus, Ivan Iossifov, Michael C. Schatz, Gholson J. Lyon

## Abstract

**Background:** INDELs, especially those disrupting protein-coding regions of the genome, have been strongly associated with human diseases. However, there are still many errors with INDEL variant calling, driven by library preparation, sequencing biases, and algorithm artifacts.

**Methods:** We characterized whole genome sequencing (WGS), whole exome sequencing (WES), and PCR-free sequencing data from the same samples to investigate the sources of INDEL errors. We also developed a classification scheme based on the coverage and composition to rank high and low quality INDEL calls. We performed a large-scale validation experiment on 600 loci, and find high-quality INDELs to have a substantially lower error rate than low quality INDELs (7% vs. 51%).

**Results:** Simulation and experimental data show that assembly based callers are significantly more sensitive and robust for detecting large INDELs (>5 bp) than alignment based callers, consistent with published data. The concordance of INDEL detection between WGS and WES is low (52%), and WGS data uniquely identifies 10.8-fold more high-quality INDELs. The validation rate for WGS-specific INDELs is also much higher than that for WES-specific INDELs (85% vs. 54%), and WES misses many large INDELs. In addition, the concordance for INDEL detection between standard WGS and PCR-free sequencing is 71%, and standard WGS data uniquely identifies 6.3-fold more low-quality INDELs. Furthermore, accurate detection with Scalpel of heterozygous INDELs requires 1.2-fold higher coverage than that for homozygous INDELs. Lastly, homopolymer A/T INDELs are a major source of low-quality INDEL calls, and they are highly enriched in the WES data.

**Conclusions:** Overall, we show that accuracy of INDEL detection with WGS is much greater than WES even in the targeted region. We calculated that 60X WGS depth of coverage from the HiSeq platform is needed to recover 95% of INDELs detected by Scalpel. While this is higher than current sequencing practice, the deeper coverage may save total project costs because of the greater accuracy and sensitivity. Finally, we investigate sources of INDEL errors (e.g. capture deficiency, PCR amplification, homopolymers) with various data that will serve as a guideline to effectively reduce INDEL errors in genome sequencing.

## Background

With the increasing use of next-generation sequencing (NGS), there is growing interest from researchers, physicians, patients and consumers to better understand the underlying genetic contributions to various conditions. For rare diseases and cancer studies, there has been increasing success with exome/genome sequencing in identifying mutations that have a large effect size for particular phenotypes [1–3]. Some groups have been trying to implement genomic and/or electronic health record approaches to interpret disease status and inform preventive medicine [4–8]. However, we are still facing practical challenges for both analytic validity and clinical utility of genomic medicine [9–13]. In addition, the genetic architecture behind most human disease remains unresolved [14–19]. Some have argued that we should bring higher standards to human-genetics research in order to return results and/or reduce false-positive reports of “causality” without rigorous standards [20, 21]. Others have reported that analytic validity for WES and WGS is still a major issue, pointing out that the accuracy and reliability of sequencing and bioinformatics analysis can and should be improved for a clinical setting [10, 11, 22–25].

There is also debate whether we should primarily in the year 2014 use whole genome sequencing (WGS) or whole exome sequencing (WES) for personal genomes. Some have suggested that a first-tier cost-effective WES might be a powerful way to dissect the genetic basis of diseases and to facilitate the accurate diagnosis of individuals with Mendelian disorders [26, 27]. Others have shown that targeted sequencing misses many things [28] and that WGS could reveal structural variants (SVs), maintains a more uniform coverage, is free of exome capture efficiency issues, and actually includes the noncoding genome, which likely has substantial importance [29–32]. Some groups directly compared WGS with WES, but thorough investigation of INDEL errors was not the focus of these comparisons [10, 23, 24, 33]. Substantial genetic variation involving INDELs in the human genome has been previously reported but accurate INDEL calling is still difficult [34–36]. There has been a dramatic decrease of sequencing cost over the past few years, and this cost is expected to decrease further with the release of the Illumina HiSeq×Ten sequencers which have capacity for nearly 18,000 whole human genomes per instrument per year. However, it is still unclear whether we can achieve a high-accuracy personal genome with a mean coverage of 30X from the Illumina HiSeq X Ten sequencers. In addition, there have been questions on the use of PCR amplification in the library preparations for NGS, although very few have characterized the PCR errors that might be complicating the detection of insertions and deletions (INDELs).

Concordance rates among INDELs detected by the GATK Unified Genotyper (v1.5), SOAPindel (v1.0) and SAMtools (v0.1.18) are reportedly low, with only 26.8% agreeing across all three pipelines [10]. Another group also reported low concordance rates for INDELs between different sequencing platforms, further showing the difficulties of accurate INDEL calling [24]. Other efforts have been made to understand the sources of variant calling errors [12]. Common INDEL issues, such as realignment errors, errors near perfect repeat regions, and an incomplete reference genome have caused problems for approaches working directly from the alignments of the reads to reference [37, 38]. De novo assembly using de Brujin graphs has been reported to tackle some of these limitations [39]. Fortunately, with the optimization of micro-assembly, these errors have been reduced with a novel algorithm, Scalpel, with substantially improved accuracy over GATK-HaplotypeCaller (v3.0), SOAP-indel (v2.01), and six other algorithms [40]. Based on validation data, the positive prediction rate (PPV) of algorithm specific INDELs was high for Scalpel (77%), but much lower for GATK HaplotypeCaller (v3.0) (45%) and SOAP-indel (v2.01) (50%) [40].

Thus, we set out to investigate the complexities of INDEL detection on Illumina reads using this most accurate INDEL-calling algorithm. Firstly, we used simulation data to understand the limits of how coverage affects INDEL calling with Illumina-like reads using GATK-UnifiedGenotyper and Scalpel. Secondly, we analyzed a dataset including high coverage WGS and WES data from two quad families (mother, father and two children), in addition to extensive high-depth validation data on an in-house sample, K8101-49685s. In order to further understand the effects of PCR amplification on INDEL calling, we also downloaded and analyzed two WGS datasets prepared with and without PCR from the well-known HapMap sample NA12878. We characterized the data in terms of read depth, coverage uniformity, base-pair composition pattern, GC contents and other sequencing features, in order to partition and quantify the INDEL errors. We were able to simultaneously identify both the false-positives and false-negatives of INDEL calling, which will be useful for population-scale experiments. We observe that homopolymer A/T INDELs are a major source of low quality INDELs and multiple signatures. As more and more groups start to use these new micro-assembly based algorithms, practical considerations for experimental design should be introduced to the community. Lastly, we explicitly address the question concerning the necessary depth of coverage for accurate INDEL calling using Scalpel for WGS on HiSeq sequencing platforms. This work provides important insights and guidelines to achieve a highly accurate INDEL call set and to improve the sequencing quality of personal genomes.

## Methods

### Analysis of Simulated Data

We simulated Illumina-like 2*101 paired-end reads with randomly distributed INDELs, which ranged from 1 bp to 100 bp. The simulated reads were mapped to human reference genome hg19 using BWA-mem (v0.7-6a) using default parameters [41]. The alignment was sorted with SAMtools (v0.1.19-44428cd) [42] and the duplicates were marked with Picard using default parameters (v1.106), resulting in a mean coverage of 93X. We down-sampled the reads with Picard to generate 19 sub-alignments. The minimum mean coverage of the sub-alignments was 4.7X and increased by 4.7X each time, before it reached the original coverage (93X). Scalpel (v0.1.1) was used as a representative of assembly based callers to assemble the reads and call INDELs from each alignment separately, resulting in 20 INDEL call-sets from these 20 alignments, using the following parameter setting: “--single --lowcov 1 --mincov 3 –outratio 0.1 --numprocs 10 --intarget”. We also used GATK-UnifiedGenotyper (v3.2-2) as a representative of alignment based callers to call INDELs from each set of alignments [43]. We followed the best practices on the GATK website, including all the pre-processing procedures, such as INDEL realignment and base recalibration. Scalpel (v0.1.1) internally left-normalized all the INDELs so we only used GATK-LeftAlignAndTrimVariants on the INDEL calls from UnifiedGenotyper. We then computed both the sensitivity and false discovery rate (FDR) for both INDEL callers, with respects to all and large (>5 bp) INDELs. The same versions and the same sets of parameter settings for BWA-mem, Picard, and Scalpel, were also used in the rest of the study, including the analysis of WGS/WES data, standard WGS and PCR-free data.

### Generation of WGS and WES data

Blood samples were collected from eight humans of two quartets from the Simons Simplex Collection (SSC). Both WGS and WES were performed on the same genomic DNA isolated from these eight blood samples. The exome capture kit used was NimbleGen SeqCap EZ Exome v2.0, which was designed to pull down 36Mb (approximately 300,000 exons) of the human genome hg19. The actual probe regions were much wider than these targeted regions, because probes also covered some flanking regions of genes, yielding a total size of 44.1Mb. All of the libraries were constructed with PCR amplification. We sequenced both sets of libraries on Illumina HiSeq2000 with average read length of 100 bp at the sequencing center of Cold Spring Harbor Laboratory (CSHL). We also generated WGS (mean coverage=30X) and WES (mean coverage=110X) data from an in-house sample K8101-49685s (not from SSC), which was extensively investigated in the later validation experiment. Exome capture for this sample was perfomed using the Agilent 44Mb SureSelect protocol and the resulting library was sequenced on Illumina HiSeq2000 with average read length of 100 bp. All of the HiSeq data from K8101-49685s have been submitted to the Sequence Read Archive (SRA) (http://www.ncbi.nlm.nih.gov/sra/) under project accession number SRX265476 (WES data) and SRX701020 (WGS data).

### Analysis of the INDELs from WGS and WES data

We excluded all of the low quality raw reads, aligned the remaining high quality ones with BWA-mem, and mark-duplicated with Picard. We used Scalpel to assemble the reads and identify INDELs under both single mode and quad mode. The single mode outputs all of the putative INDELs per person, and the quad mode outputs only the putative de novo INDELs in the children in a family. We expanded each of the exons by 20 bp upstream and 20 bp downstream in order to cover the splicing sites and we called this set of expanded regions the “exonic targeted regions”. The exonic targeted regions are fully covered by the exome capture probe regions. We excluded INDELs that were outside the exonic targeted regions in the downstream analysis.

We left-normalized the INDELs and compared the two call sets for the same person using two criteria: exact-match and position-match. Position-match means two INDELs have the same genomic coordinate, while exact-match additionally requires that two INDELs also have the same base-pair change(s). We called the INDELs in the intersection based on exact-match as WGS-WES intersection INDELs. Further, we named the INDELs only called from one dataset as “WGS-specific” and “WES-specific” INDELs, respectively. Regions of the above three categories of INDELs were partitioned and investigated separately. In particular, we focused on regions containing short tandem repeats (STR) and homopolymers. We used BedTools (v2.18.1) with the region file from lobSTR (v2.04) to identify homopolymeric regions and other STR (dual repeats, triplets and etc.) in the human genome [44–46].

### Generating summary statistics of alignment from WGS and WES

We used Qualimap (v0.8.1) to generate summary statistics of the alignment files of interest [47]. For a certain region, we define the proportion of a region covered with at least X reads to be the coverage fraction at X reads. In addition to the coverage histograms, we also computed the coefficient of variation (C_v_) to better understand the coverage uniformity of the sequencing reads. An unbiased estimator of C_v_ can be computed by 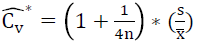, where s represents the sample standard deviation and 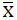 represents the sample mean. In our case, 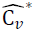 asymtotically approaches to 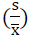 as the sample size (n) of the data is usually greater than 10,000. The reference genome used here is hg19. There were four region files that we used for this part of the analysis. The first one is the exon region bed file from NimbleGen. We generated the other three region files by expanding 25 bp upstream and downstream around loci of WGS-WES intersection INDELs, WGS-specific INDELs, and WES-specific INDELs, respectively. We followed all of the default settings in Qualimap except for requiring the homopolymer size to be at least five (-hm 5). Finally, we used Matplotlib to generate the figures with the raw data from Qualimap under the Python environment 2.7.2 [48].

### Generation of MiSeq validation data of sample K8101-49685s

We randomly selected 200 INDELs for validation on an in-house sample K8101-49685s from each of the following categories: 1) INDELs called from both WGS and WES data (WGS-WES intersection), 2) WGS-specific INDELs, 3) WES-specific INDELs. Out of these 600 INDELs, 97 were covered with more than 1,000 reads in the previous MiSeq data set reported by Narzisi *et al*. Hence, we only performed additional Miseq validation on the remaining 503 loci [40]. PCR primers were designed using Primer 3 to produce amplicons ranging in size of 200 - 350 bp, with INDELs of interest located approximately in the center. Primers were obtained from Sigma-Aldrich in 96-well mixed-plate format, 10 µmol/L dilution in Tris per oligonucleotide. 25 µL PCR reactions were set up to amplify each INDEL of interest using K8101-49685s’ genomic DNA as template and LongAmp Taq DNA polymerase (New England Biolabs). PCR products were visually inspected for amplification efficiency using 1.5% agarose gel electrophoresis, and then pooled for ExoSAP-IT (Affymetrix) cleanup. The cleanup product was purified using QIAquick PCR Purification Kit (Qiagen) and quantified by Qubit dsDNA BR Assay Kit (Invitrogen). Subsequently, a library construction was performed following the TruSeq Nano DNA Sample Preparation Guide for the MiSeq Personal Sequencer platform (Illumina). Before loading onto the MiSeq machine, the quality and quantity of the sample was reevaluated using the Agilent DNA 1000 Kit on the Agilent Bioanalyzer and with quantitative PCR (Kapa Biosystems).

We generated high quality 250 bp paired-end reads with an average coverage of 55,000X over the selected INDELs. We aligned the reads with BWA-MEM (v0.7.5a) to hg19, sorted the alignment with SAMtools (v0.1.18) and marked PCR duplicates with Picard (v1.91). The alignment quality control showed that 371 out of the 503 loci were covered with at least 1,000 reads in the data and we only considered these loci in the downstream analysis. Therefore, we have validation data on 160, 145 and 161 loci from the WGS-WES intersection, WGS-specific, and WES-specific INDELs, respectively. As reported by Narzisi *et al*, mapping the reads containing a large INDEL (near or greater than half the size of the read length) is problematic. This was particularly difficult when the INDEL is located toward either end of a read [40]. To avoid this, we used very sensitive settings with Bowtie2 (--end-to-end --very-sensitive --score-min L,-0.6,-0.6 --rdg 8,1 --rfg 8,1 --mp 20,20) to align the reads because it can perform end-to-end alignment and search for alignments with all of the read characters [49]. We generated the true INDEL call set by two steps: 1) used GATK UnifiedGenotyper to call INDELs from the BWA-MEM alignment, 2) performed manual inspection on the large INDELs from the Bowtie2 alignment (require at least 25% of the reads supporting an INDEL) [43]. The alignments were realigned with the GATK (v2.6-4) IndelRealigner and base quality scores were recalibrated before variants were called with UnifiedGenotyper. Left-normalization was performed to avoid different represenations of a variant. An INDEL was considered valid if a mutation with the same genomic coordinate and the same type of variation exists in the validation data. For example, an insertion call would not be considered valid if the variant with the same coordinate in the validation data was instead a deletion. All of the MiSeq data can be downloaded from the Sequence Read Archive under project accession number SRX386284 (Accession number: SRR1575211, SRR1575206, SRR1042010).

### Classifications of INDEL with calling quality based on the validation data

We previously benchmarked Scalpel with respect to the coverage of the alternative allele 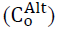 and the k-mer Chi-Square scores (χ^2^). Scalpel applied the standard formula for the Chi-Square statistics and applied to the K-mer coverage of both alleles of an INDEL.

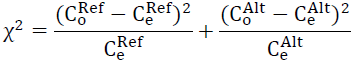

where 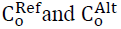 are the observed k-mer coverage for the reference and alternative alleles, 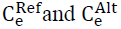 are the expected k-mer coverage, i.e. 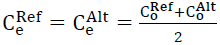.

Here we used 466 INDELs from the validation data to understand the relationship between the FDR and these two metrics (Supplemental Figure S4). Our validation data showed that with the same χ^2^, INDELs with a lower 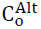 tend to have a higher FDR, especially for INDELs with 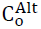 not greater than 10 (Supplemental Figure S4). For INDELs with relatively the same 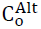, a higher χ^2^ also made them less likely to be valid. We noticed that the calling quality could be determined by the error rate inferred by these two metrics. To achieve a consistent accuracy for INDELs with different 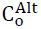, we classified INDEL calls and determined the calling quality with the below criteria: High quality INDELs: low error-rate (7%) INDELs meeting any of the three cutoffs: 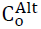 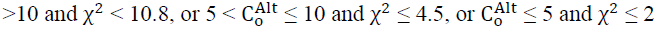; Low quality INDELs: high error-rate (51%) INDELs meeting the following cutoff: 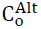 ≤ 10 and χ^2^ > 10.8; Moderate quality: The remaining INDELs that do not fall into the above two categories.

### Analysis of PCR-free and standard WGS data of NA12878

We downloaded PCR-free WGS data of NA12878 (access Code: ERR194147), which is publicly available in the Illumina Platinum Genomes project. We also download another WGS dataset of NA12878 with PCR amplification during library prepartion, and we called it standard WGS data (SRA access Code: SRR533281, SRR533965, SRR539965, SRR539956, SRR539947, SRR539374, SRR539357). Both data were generated on the Illumina HiSeq 2000 platform. Although the PCR-free data was not supposed to have any PCR duplicates, we observed a duplication rate of 2% as reported by Picard, and we excluded these reads, yielding 50X mean coverage for both datasets after removing PCR duplicates. We used the same methods for alignment, INDEL calling, and downstream analysis as described above. INDELs outside the exonic targeted regions were not considered in the downstream analysis.

### Analysis of INDEL detection sensitivity in WGS data

We were interested to know how depth of coverage affects the sensitivity of INDEL detection in WGS data. To accurately measure this sensitivity, one needs a robust call set as a truth set. Fortunately, we had exact-match INDELs concordant between high coverage WGS and high coverage WES data. We therefore measured sensitivity based on these WGS-WES intersection INDELs, rather than on the whole set of INDELs, which might contain more false positives. We down-sampled each WGS dataset to mean coverages of 20X, 32X, 45X and 57X. We then used Scalpel to call INDELs from the resulting 4 sub-alignment files for each sample and computed the sensitivity at a certain mean coverage (X) for each sample by the equation:

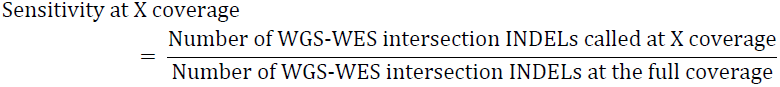

This equation measures how many of the WGS-WES intersection INDELs can be discovered as a function of read depth. We also analyzed the WGS-WES intersection INDEL call set in terms of zygosity: WGS-WES intersection heterozygous and homozygous INDEL, subsequently measuring the sensitivity with respect to different zygosities.

## Results and discussion

### Simulated data: Characterizing alignment and assembly based callers at different coverage

We started our study with asking whether depth of sequencing coverage affect different kinds of INDEL calling algorithms. (e.g. assembly based callers and alignment based callers). Thus, we began with simulated reads with known error rates across the genome to answer this question. We used GATK-UnifiedGenotyper (v3.2-2) and Scalpel (v0.1.1) as a representative of alignment based callers and assembly based callers, repsectively. Figure 1A shows that for both algorithms, higher coverage improves sensitivity of detecting both general INDELs (i.e. any size starting from 1 bp) and large INDELs (i.e. size greater than 5 bp). For general INDEL detection with both algorithms, this improvement did not saturate until a mean coverage of 28X. Furthermore, detecting large INDELs was more difficult than general INDELs because the increase of sensitivity did not saturate until reaching a mean coverage of 42X. However, there were substantial differences of sensivity performance between these two algorithms for large INDEL detection. We noticed that even at a very high coverage (mean coverage = 90X), GATK-UnifiedGenotyper could call only about 52% of the large INDELs while Scalpel could reveal more than 90% of them. This is because GATK-UnifiedGenotyper tries to infer genotypes from alignment and large INDELs could complicate or distort the correct mapping. To achieve a sensitivity of 90% with Scalpel, a mean coverage of 30X was required for general INDEL detection while 90X was needed to detect large INDELs at a similar sensitivity. This showed that much higher coverage is needed for large INDEL detection, especially to maintain coverage across the INDEL and to have enough partially mapping or soft-clipped reads to use for the micro-assembly.

**Figure 1.**
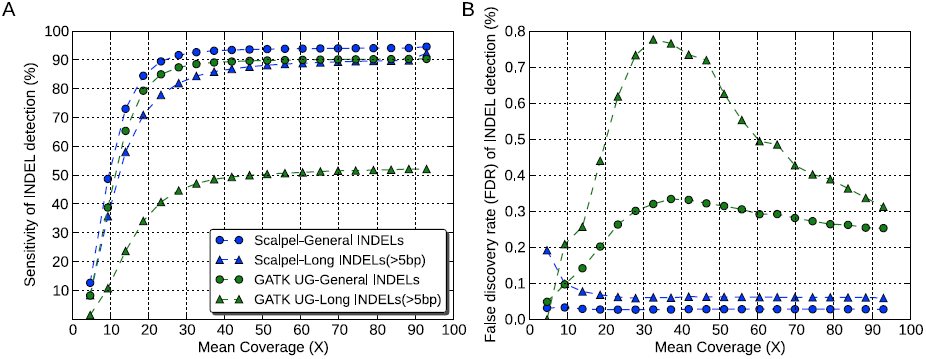
Performance comparison between the Scalpel and GATK-UnifiedGenotyper in terms of sensitivity (A) and false discovery rate (B) at different coverage based on simulation data. Each dot represent one down-sampled experiment. Round dots represent performance of general INDELs (i.e. INDELs of size starting at 1 bp) and triangles represent performance of large INDELs (i.e. INDELs of size greater than 5 bp). The data of Scalpel was shown in blue while GATK-UnifiedGenotyper was shown in green.

The FDRs of Scalpel were robust to the changes in coverage while GATK-UnifiedGenotyper’s FDRs were affected by coverage. For the detection of large INDELs with Scalpel, the FDRs marginally decreased as the mean coverage increased from 5X to 28X, and remained basically the same again from 33X to 93X (Figure 1B). This indicates that for large INDELs, insufficient coverage results in more assembly errors, which results in a higher error rate for micro-assembly variant calling. Based on the simulation data, a mean coverage of at least 30X is needed to maintain a reasonable FDR for Scalpel. In contrast, FDRs of GATK-UnifiedGenotyper are much higher and more unstable at different coverages, especially for large INDELs. Nonetheless, since these results were based on simulation data, which does not include the effects of any sequencing artifacts on INDEL calling, these values establish the upper bound of accuracy and performance compared to genuine sequence data. Previous studies reported that local assembly allows to call INDELs much larger than those that can be identified by the alignment [13, 40, 50]. Consistent with previous reports, our simulated data suggested that assembly based callers can reveal a much larger spectrum of INDELs than alignment based callers, in terms of their size. Furthermore, Narzisi *et al.* recently reported that Scalpel is more accurate than GATK-HaplotypeCaller and SOAPindel, especially within regions containing near-perfect repeats [40]. Thus, in order to control for artifacts from callers, we chose to use Scalpel as the only INDEL caller in our downstream analysis on the experimental data, which could help to better clarify differences between data types.

### WGS vs. WES: Low concordance on INDEL calling

We analyzed a dataset including high coverage WGS and WES data from eight samples in the SSC. To make a fair comparison, the INDEL calls were only made from the exonic targeted regions as explained in the Methods. The mean INDEL concordance between WGS and WES data was low, 53% using exact-match and 55% using position-match (Figure 2, Table 1). Position-match means the two INDELs have the same genomic coordinate, while exact-match additionally requires that the two INDELs also have the same base-pair change(s) (see Methods). When we excluded regions with less than one read in either dataset, the mean concordance rates based on exact match and position-match increased to 62% and 66%, respectively (Table 1). If we excluded regions with base coverage in either dataset with less than 20, 40, 60, or 80 reads, the mean concordance rate based on exact-match and position-match both continued to increase until reaching a base coverage of 80 reads (Table 1). This showed that some INDELs were missing in either dataset because of low sequencing efficiency in those regions. Although WES data had higher mean coverage than WGS data, we were surprised to see that in regions requiring at least 80 reads, there were more INDELs that were specific to WGS data than WES data (21% vs. 4%). Regions with excessive coverage might indicate problems of sequencing or library preparation, and this highlights the importance of coverage uniformity in WGS (Figure 3A & B, Table 2). It should be noted that mapping artifacts could also be a possible reason. For example, the reads may originate in regions which are absent from the reference genome, such as copy number variants [51]. Based on exact-match, the proportion of the WGS-specific INDELs was 2.5-fold higher than that of WES-specific INDELs (34% vs. 14%). This difference was even larger based on position-match (3-fold). In principle, the reasons for this could be either high sensitivity of INDEL detection with WGS data or high specificity of INDEL detection with WES data, and we will examine these options in more detail below.

**Figure 2.**
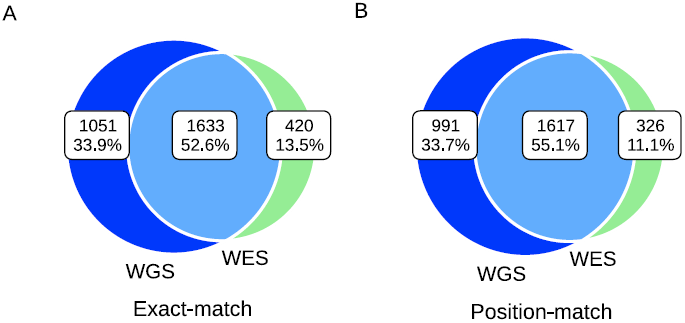
Mean concordance of INDELs over eight samples between WGS (blue) and WES (green) data. Venn diagram showing the numbers and percentage of shared between data types based on (A) Exact-match (B) Position-match. The mean concordance rate increased when we required at least a certain number of reads in both data (Table 1).

**Figure 3.**
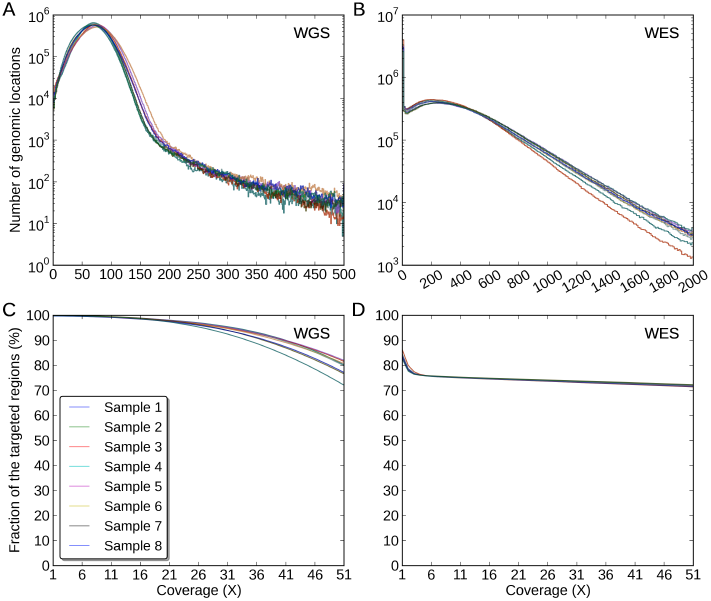
Coverage distributions of the exonic targeted regions in (A) the WGS data, (B) the WES data. The Y-axis for A) and B) is of log10-scale. The coverage fractions of the exonic targeted regions from 1X to 51X in (C) the WGS data, (D) the WES data.

**Table 1.**
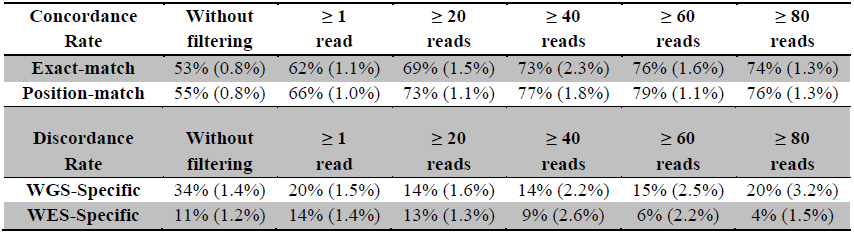
**Mean concordance and discordance rates of INDEL detection between WGS and WES data in different regions**. The data is shown in the following order: 1) regions without filtering, and regions filtered by requiring base coverage to be at least 2) one read, 3) 20 reads, 4) 40 reads, 5) 60 reads, or 6) 80 reads in both data. The mean discordance rate is calculated based on position-match, which is the percentage of INDELs specific to either dataset. The standard deviation is shown in parenthesis.

**Table 2.** **Mean coefficients of variation of coverage with respects to the following regions: WGS-WES intersection INDELs, WGS-specfic INDELs, and WES-specific INDELs**. WGS-WES intersection INDELs means the INDELs called from both WGS and WES data. WGS-specific INDELs means the INDELs only called from the WGS data. The standard deviation is shown in parenthesis.

|  | Exonic targeted regions | WGS-WES intersection INDEL regions | WGS-specific INDEL regions | WES-specific INDEL regions |
| --- | --- | --- | --- | --- |
| WGS | 39.4% (1.9%) | 47.2% (3.0%) | 75.3% (5.7%) | 56.1% (9.6%) |
| WES | 109.3% (1.5%) | 96.8% (3.2%) | 281.5% (13.3%) | 117.4% (22.8%) |

### Coverage distributions of different regions in WGS and WES data

An ideal sequencing experiment should result in a high number of reads covering a region of interest uniformly. Using the eight SSC samples, we investigated the coverage behaviours of the WGS and WES data by the following: distribution of the read depth, mean coverage, coverage fraction at X reads, coefficient of variation (C*_v_*) (See methods). Hence, ideally one should expect to see a normal distribution of read depth with a high mean coverage and a small C*_v_*. Comparisons of the coverage distributions are shown in the following order: 1) Exonic targeted regions, i.e. the exons that the exome capture kit was designed to pull down and enrich; 2) WGS-WES intersection INDEL regions, i.e. the regions where WGS and WES revealed the identical INDELs based on exact-match; 3) WGS-specific INDEL regions, i.e. the regions where only WGS revealed INDELs based on position-match; 4) WES-specific INDEL regions, i.e. the regions where only WES revealed INDELs based on position-match.

Firstly, in the exonic targeted regions, the mean coverages across eight samples were 71X and 337X for WGS and WES data, respectively (Figure 3A & B, Supplemental Table S1). We noticed that there was a recovery issue with WES in some regions, as the coverage fraction at 1X was 99.9% in WGS data but only 84% in WES data, meaning that 16% of the exonic targeted regions were not recovered, which could be due to capture inefficiency or other issues involving DNA handling during the exome library preparation and sequencing protocols (Figure 3C & D, Supplemental Table S2). The coverage was much more uniform in the WGS data than that in the WES data because C*_v_* of the WGS data was much lower (39% vs. 109%, Figure 3A & B, Table 2). Secondly, in the WGS-WES intersection INDEL regions, the mean coverage across eight samples were 58X and 252X for WGS and WES data, respectively (Supplemental Figure S1A & B, Supplemental Table S1). We noticed that there was an increase of coverage uniformity for WES in the WGS-WES intersection INDEL regions, relative to the exonic targeted regions, because C*_v_* was lower (109% vs. 97%) (Table 2, Figure 3B, Supplemental Figure S1b). We noticed WGS was able to reveal WGS-WES intersection INDELs at a much lower coverage relative to WES, which we attribute to a better uniformity of reads across the genome (C*_v_*: 47% vs. 97%, Table 2, Supplemental Figure S1A & B). The coverage distributions were skewed in the WES data, with some regions poorly covered and other regions over saturated with redudant reads.

Thirdly, in WGS-specific INDEL regions, the mean coverages across eight samples were 61X and 137X for WGS and WES data, respectively (Figure 4, Supplemental Table S1). Compared to the entire exonic targeted regions, the mean coverage for WES data was significantly reduced in these regions (137X vs. 337X), and 44% of the regions were not covered with a single read (Figure 4, Supplemental Table S2). We noticed that compared to the WGS data, the WES data poorly covered these regions with 20 reads or more (94% vs. 31%, Figure 4C & D). In these regions, the coverage unifomity of the WES data was much lower than that of the WGS data (C*_v_*: 282% vs. 75%, Figure 4A & B, Table 2). The reason why WES data missed these INDELs could be insufficient coverage around the INDELs in these regions. Finally, in WES-specific INDELs regions, the mean coverages across eight samples were 41X and 172X for WGS and WES data, respectively (Supplemental Figure S2A & B, Supplemental Table S1). In these regions, both data had a relatively high coverage and the WES data covered most these regions with at least one read (Supplemental Figure S2C & D). However, we noticed that the WES data still had a much lower coverage unifomity (C*_v_*: 117% vs. 56%, Table 2). In order to better understand these issues, we used the WGS-WES intersection INDEL set as a positive control and proceeded to assess each call set with newly developed quality criteria.

**Figure 4.**
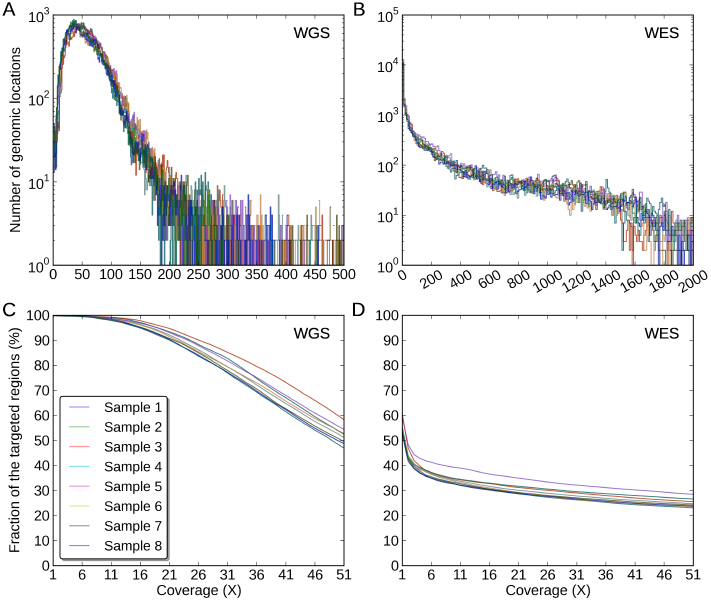
Coverage distributions of the WGS-specific INDELs regions in (A) the WGS data, (B) the WES data. The Y-axis for A) and B) is of log10-scale. The coverage fractions of the WGS-specific INDELs regions from 1X to 51X in (C) the WGS data, (D) the WES data.

### MiSeq validation of INDELs in WGS and WES data on the sample K8101-49685s

In order to understand error rates and behaviours of the INDEL call from the WGS and WES data, we randomly selected 200 INDELs for MiSeq validation on the sample K8101-49685s from each of the following categories: 1) INDELs called from both WGS and WES data (WGS-WES intersection INDELs), 2) WGS-specific INDELs, 3) WES specific INDELs. First, the validation rate of WGS-WES intersection INDELs was in fact very high (95%), indicating INDELs called from both WGS and WES data were mostly true-positives (Table 3). Second, the validation rate of WGS-specific INDELs was much higher than that of WES-specific INDELs (84% vs. 57%). Third, among the validation set, large INDELs (> 5 bp) that were called from both the WGS and WES data were 100% valid, while the validation rate of large INDELs that were specific to the WGS data was only 76%. However, we noticed that there was only one large INDEL specific to the WES data that we selected for validation. Since the sampling was performed randomly, we examined the original call set to understand this phenomenon. Only 9% of the WGS-WES intersection INDELs (176) and 21% of the WGS-specific INDELs (106) were greater than 5 bp (Table 4). But we were surprised to see that only 1.5% of the WES-specific INDELs were greater than 5 bp, meaning only 10 INDELs were large according to our definition. This showed that the WES data missed most large INDELs, which we speculate might be due to capture deficiency or some other procedure related to the process of exome capture and sequencing. In particular, large INDELs could disrupt the base pairing that occurs during the exome capture procedure, which would then result in insufficient coverage in those regions (Figure 4).

**Table 3.**
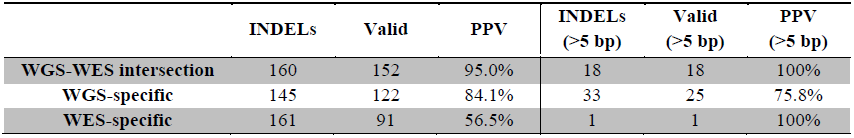
**Validation rates of WGS-WES intersection INDELs, WGS-specfic, and WES-specific INDELs.** We also calculated the validation rates of large INDELs (>5 bp) in each category. The validation rate, positive predictive value (PPV), is computed by the following: PPV=#TP/(#TP+#FP), where #TP is the number of true-positive calls and #FP is the number of false-positive calls.

**Table 4.** **Number and fraction of large INDELs in the following INDEL categories: 1)** WGS-WES intersection INDELs**, 2) WGS-specific, and WES-specific**.

|  | **All**<br>**INDELs** | **Large INDELs**<br>**(>5 bp)** | **Fraction of large**<br>**INDELs (>5 bp)** |
| --- | --- | --- | --- |
| **WGS-WES intersection** | 2009 | 176 | 8.8% |
| **WGS-specific** | 494 | 104 | 21.1% |
| **WES-specific** | 674 | 10 | 1.5% |

### Assessment of the INDEL calls sets from WGS and WES

To understand the error profile of the WGS and WES data with a larger sample size, we developed a classification scheme based on the validation data and applied them to the eight samples in the Simons Simplex Collection (SSC). Three combinations of thresholds were used to define the calling quality of an INDEL call as either high, moderate or low quality based on the following two metrics: the coverage of the alternative allele and the k-mer Chi-Square score of an INDEL (see Methods). Based on those cutoffs, there was 7.3-fold difference between high-quality and low-quality INDELs in terms of their error rates (7% vs. 51%). This suggests that our classification scheme is able to effectively distinguish behaviours of problematic INDEL calls from likely true-positives. Our classification scheme is also useful for eliminating false *de novo* INDEL calls in family-based studies (see Supplemental Note 1). Futhermore, WGS-WES intersection and WGS-specific INDELs seem to be reliable calls, and the majority of the INDELs in these two call sets were of high-quality, 89% and 78% respectively. Only a very small fraction of them were of low-quality, 2% and 7% respectively. (Figure 5, Supplemental Table S3). In contrast, for WES-specific INDELs, there was a striking enrichment of low-quality events (41%), and a 4.1-fold decrease of the high-quality events (22%). Notably, among these 8 samples. there were 991 WGS-specific INDELs and 326 WES-specific INDELs, and from these, 769 of WGS-specific INDELs and 71 of the WES-specific INDELs were of high quality. This comparison determined that WGS yielded 10.8-fold more high quality INDELs than WES according to our classification scheme. Futhermore, WES produced 133 low quality INDELs per sample, while WGS only produced 71 low quality INDELs per sample. That being said, WES yielded 1.9-fold more low quality INDELs. This indicates WES tends to produce a larger fraction of error-prone INDELs, while WGS reveals a more sensitive and specific set of INDELs.

**Figure 5.**
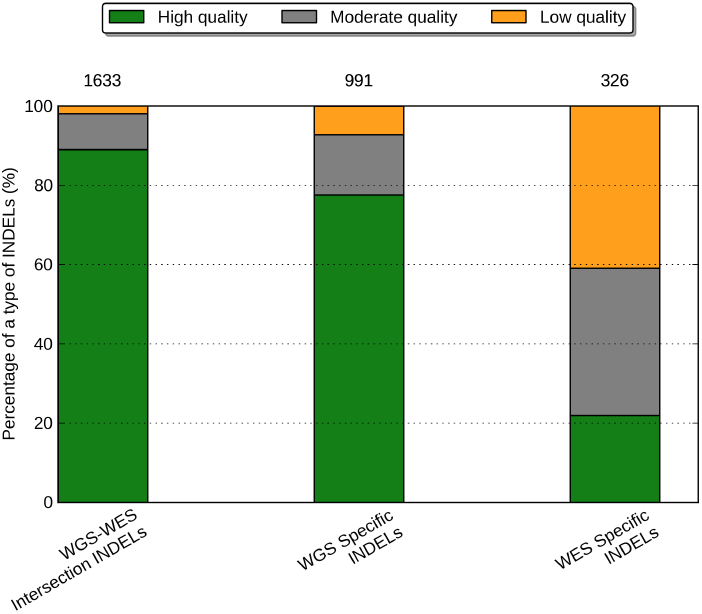
Percentage of high quality, moderate quality and low quality INDELs in three call set. (A) the WGS-WES intersection INDELs, (B) the WGS-specific INDELs, (C) the WES-specific INDELs. The numbers on top of a call set represent the mean number of INDELs in that call set over eight samples.

In order to understand what was driving the error rates in different data sets, we partitioned the INDELs according to their sequence composition: homopolymer A (poly-A), homopolymer C (poly-C), homopolymer G (poly-G), homopolymer T (poly-T), short tandem repeats (STR) except homopolymers (other STR), and non-STR. We noticed that for the high quality events, the majority of the WGS-WES intersection INDELs (70%) and WGS-specific INDELs (67%) were within non-STR regions (Figure 6, Supplemental Table S4 & S5). On the contrary, the majority of the high quality INDELs specific to WES were within poly-A (24%) and poly-T regions (30%). When we compared the low quality INDELs to the high quality INDELs, there were consistent enrichment of homopolymer A or T (poly-A/T) INDELs in all three call sets, 2.3-fold for WGS-WES intersection events, 2.1-fold for WGS-specific events, and 1.5-fold for WES-specific events. The WES-specific call set contained a much higher proportion (83%) of Poly-A/T INDELs from the low-quality INDELs, relative to the WGS-WES intersection call set (44%), and the WGS-specific call set (45%). This suggested that poly-A/T is a major contributor to the low quality INDELs, which gives rise to much more INDEL errors. We explored this further in the comparison of PCR-free and standard WGS data below.

**Figure 6.**
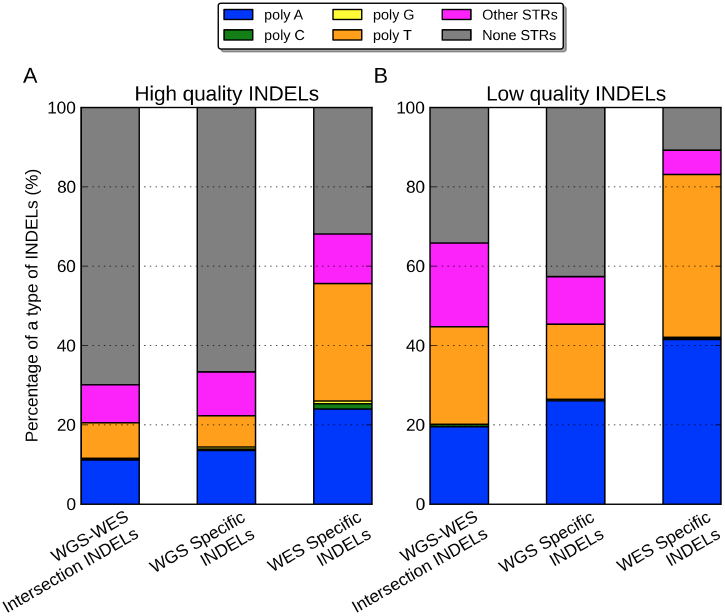
Percentage of poly-A, poly-C, poly-G, poly-T, other-STR, and non-STR in three call set. (A) high quality INDELs, (B) low quality INDELs. In both figures, from left to the right are WGS-WES intersection INDELs, WGS-specific INDELs, and WES-specific INDELs.

### Sources of multiple signatures in WGS and WES data

Another way of understanding INDEL errors is to look at multiple signatures at the same genomic location. Multiple signatures means that for the same genomic location, there are more than one INDELs called. If we assume only one signature can be the true INDEL in the genome, any additional signatures would represent false-positive calls. So if we have a higher number of multiple signatures, it means that these reads contained more INDEL errors or the algorithm tends to make more mistakes in these regions. We combined the call sets from both datasets and identified multiple signatures in the union set for each sample. In order to understand the error behaviors in the above assessment, we also partitioned the signatures by the same regional criteria. We noticed that the poly-A/T INDELs are the major source of multiple signatures, which are enriched in WES data (72% for WES vs. 54% for WGS). In particular, there is a higher number of poly-A (35 vs. 25) and poly-T (36 vs. 16) INDEL errors in the WES data than in the WGS data (Figure 7, Supplemental Table S6).

**Figure 7.**
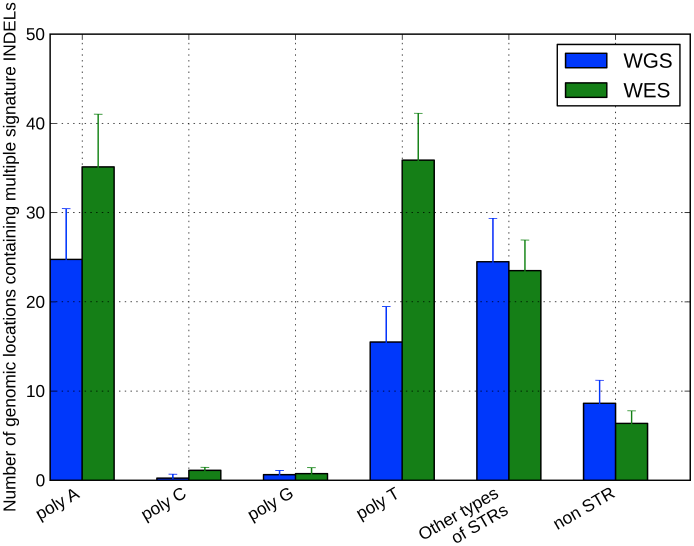
Numbers of genomic locations containing multiple signature INDELs in WGS (blue) and WES data (green). The height of the bar represents the mean across eight samples and the error bar represent the standard deviation across eight samples.

We investigated the source of multiple signatures by the numbers of reads containing homopolymer INDELs inferred by the CIGAR code (Figure 8). Figure 8 showed that there is a much higher proportion of poly-A/T INDELs in the WES-specific regions from both WGS (56%) and WES data (64%), relative to other regions. In addition, WES data has also 6.3-fold more reads than WGS data in the regions with INDELs specific to WES data (11251 vs. 1775, Supplemental Table S7). According to Qualimap, a large number of homopolymer indels might indicate a problem in sequencing for that region. Here we particularly identified the effects of these problematic sequencing reads on INDEL calling, which revealed more multiple signatures of poly-A/T INDELs.

**Figure 8.**
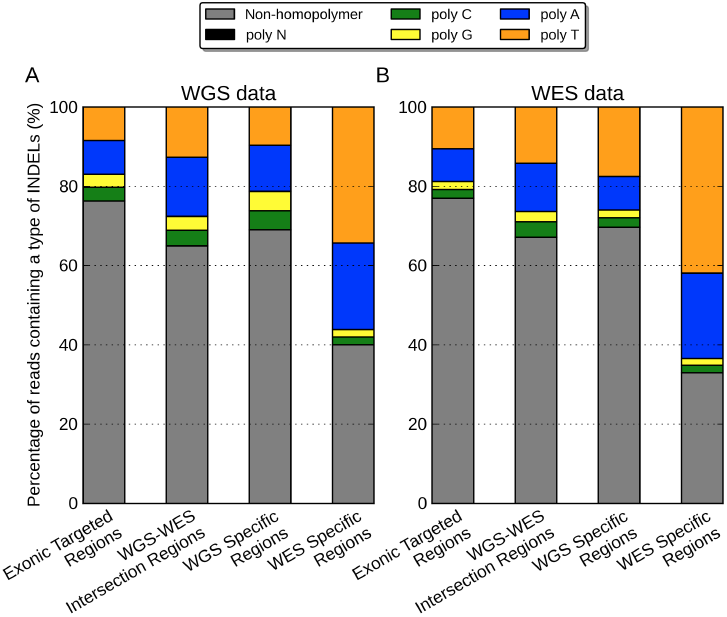
Percentage of reads near regions of Non-homopolymer, poly-N, poly-A, poly-C, poly-G, poly-T in (A) WGS data, (B) WES data. In both figures, from left to the right are exonic targeted regions, WGS-WES intersection INDELs, WGS-specific INDELs, and WES-specific INDELs.

### Standard WGS vs. PCR-free: assessment of INDELs calling quality

The concordance rate within the exonic targeted regions between standard WGS (defined as WGS involving PCR during library construction) and PCR-free data on NA12878 using exact-match and position-match were 71% and 76%, respectively (Figure 9). Note that both data used here are WGS data, so it is not surprising that these concordance rates were higher than those between WGS and WES, even for regions having at least one read in both datasets. Based on exact-match, the proportion of INDELs specific to standard WGS data was 18%, which is 1.6-fold higher than the proportion of INDELs specific to PCR-free data (11%). This ratio was similar based on position-match (1.7-fold). Like previous assessments, we classified the three call sets with respect to calling quality. We again used the INDELs called from both standard WGS and PCR-free data as a positive control. Figure 10 shows that 89% of the standard WGS & PCR-free intersection INDELs are considered as high quality, 9% as moderate quality, and only 2% as low quality. However, for INDELs specific to standard WGS data, there is a large proportion of low quality events (61%), and a very limited proportion are of high quality (7%). There were on average 310 INDELs specific to PCR-free data and 538 INDELs specific to standard WGS data. Notably, 177 of the PCR-free-specific INDELs and 40 of the standard-WGS-specific INDELs were of high quality, suggesting that in these specific regions, PCR-free data yielded 4.4-fold more high quality INDELs than standard WGS data. Furthermore, 326 of the standard-WGS-specific INDELs were of low quality, while in the PCR-free-specific call set, 52 INDELs were of low quality. That being said, in regions specific to data types, standard WGS data yielded 6.3-fold more low quality INDELs. Consistent with the comparisons between WGS and WES data, this suggested PCR amplification induced a large number of error-prone INDELs to the library, and we could effectively increase INDEL calling quality by reducing the rate of PCR amplification.

**Figure 9.**
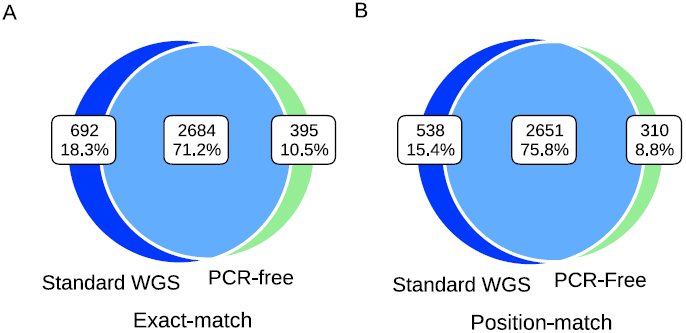
Concordance of INDEL detection between PCR-free and standard WGS data on NA12878. Venn diagram showing the numbers and percentage of shared between data types based on (A) Exact-match (B) Position-match.

**Figure 10.**
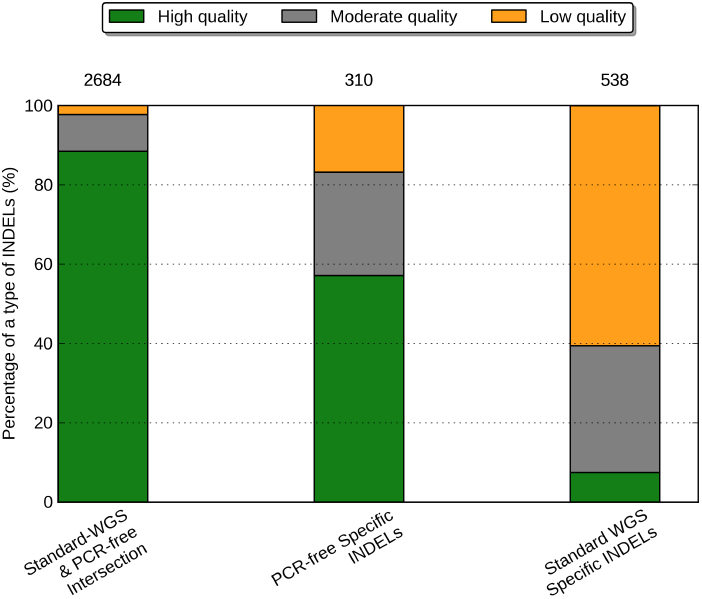
Percentage of high quality, moderate quality and low quality INDELs in two datasets. (A) the PCR-free & standard WGS INDELs, (B) the PCR-free-specific INDELs, (C) the standard-WGS-specific INDELs. The numbers on top of a call set represent the number of INDELs in that call set.

To understand the behaviors of errors in the poly-A/T regions, we partitioned the INDEL call set by the same six regions again. We noticed that for the high quality events, a majority of the standard WGS & PCR-free intersection INDELs (68%) were within non-STR regions (Figure 11). The proportion of poly-A/T INDELs was small for the standard WGS & PCR-free intersection call set (20%), larger for PCR-free-specific call set (35%), and even larger for standard-WGS-specific call set (51%). This was similar to the WGS and WES comparisons because there would be more poly-A/T INDELs when a higher rate of PCR amplification was performed. A majority of the high quality INDELs specific to standard WGS data were within poly-A (24%) and poly-T regions (38%). When we compared the low quality INDELs to the high quality ones, there was consistent enrichment of poly-A/T INDELs in all three call sets, 2.3-fold for standard WGS & PCR-free intersection events, 2.3-fold for PCR-free-specific events, and 1.3-fold for standard-WGS-specific events. For INDELs specific to standard WGS data and PCR-free data, poly-A/T INDELs represented a large proportion of the low quality INDELs: 80% and 62%, respectively. Ross *et al.* previously reported that for human samples, PCR-free library construction could increase the relative coverage for high AT regions from 0.52 to 0.82, resulting in a more uniform coverage [22]. This again suggested that PCR amplification could be a major source of low quality poly-A/T INDELs, and a PCR-free library construction protocol might be one possible solution to improve the accuracy of INDEL calls.

**Figure 11.**
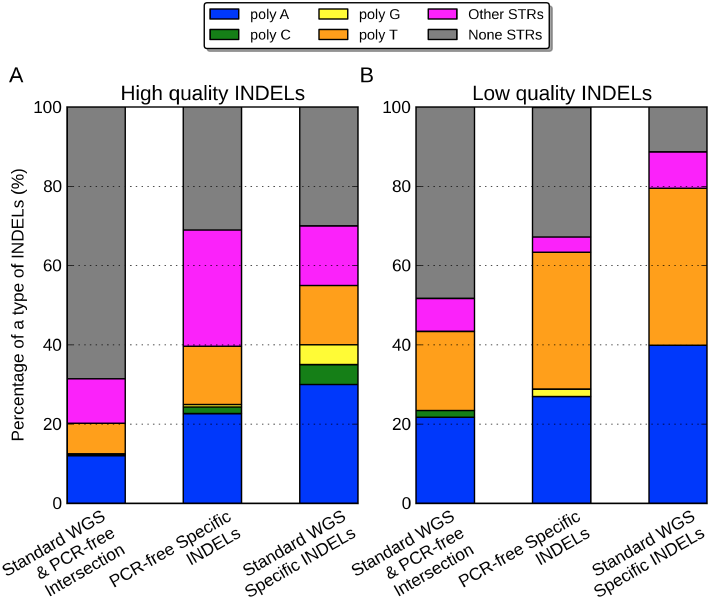
Percentage of poly-A, poly-C, poly-G, poly-T, other-STR, and non-STR in (A) high quality INDELs, (B) low quality INDELs. In both figures, from left to the right are PCR-free & standard WGS INDELs, INDELs specific to PCR-free data, and INDELs specific to standard WGS data.

### What coverage is required for accurate INDEL calling?

Ajay *et al.* 2011 reported that the number of SNVs detected exponentially increased until saturation at 40-45X average coverage [52]. However, it was not clear what the coverage requirement should be for INDEL detection. To answer this question, we down-sampled the reads, called INDELs again, and measured corresponding sensitivity for each sample using the WGS-WES intersection calls as our truth set (Methods). Figure 12A shows that we are missing 25% of the WGS-WES intersection INDELs at a mean coverage of 30X. Even at 40X coverage recommended by Ajay *et al.* 2011 [52], we could only discover 85% of the WGS-WES intersection INDELs. We calculated that WGS at 60X mean coverage (after removing PCR duplicates) from the HiSeq 2000 platform is needed to recover 95% of INDELs with Scalpel, which is much higher than current sequencing practice (Figure 12A). If economically possible, WGS at 60X mean coverage with PCR-free library preparation would generare even more ideal sequencing data for INDEL detection.

**Figure 12.**
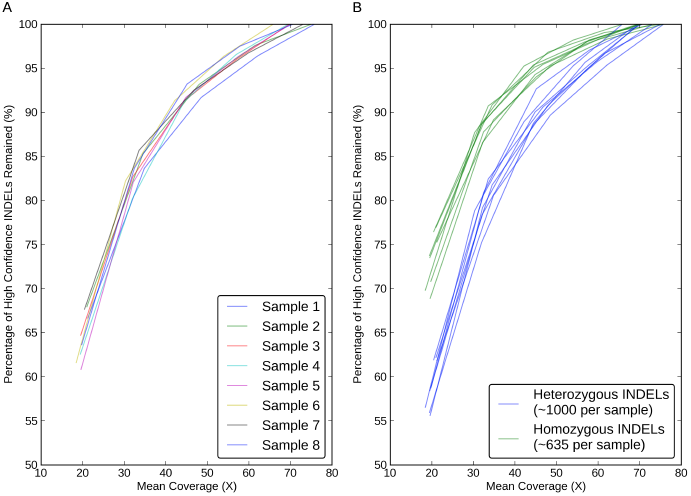
Sensitivity performance of INDEL detection with eight WGS datasets at different mean coverages on Illumina HiSeq2000 platform. The Y-axis represents the percentage of the WGS-WES intersection INDELs revealed at a certain lower mean coverage. (A) Sensitivity performance of INDEL detection with respects with each sample, (B) Sensitivity performance of heterozygous (blue) and homozygous (green) INDEL detection were shown seperately.

Some groups previously reported that determining heterozygous SNPs requires higher coverage than homozygous ones [53]. The sensitivity of heterozygous SNP detection was limited by depth of coverage, which requires at least one read from each allele at any one site and in practice much more than one read to account for sequencing errors [54]. However, the read depth requirement of INDEL detection in terms of zygosity has not been well understood. To answer this question, we took the WGS-WES intersection INDELs and partitioned them by zygosities. We first plotted the pair-wise coverage relationship between WGS and WES for each WGS-WES intersection INDEL. Supplemental Figure S3 shows that the detection of homozygous INDELs starts with a lower coverage, which is consistent in both WGS and WES datasets, although the rest of the homozygotes and heterozygotes were highly overlapping. To further understand this phenomenon, we measured the sensitivity again for heterozygous INDELs and homozygous INDELs separately. At a mean coverage of 20X, the false negative rates of WGS-WES intersection INDELs was 45% for heterozygous INDELs and 30% for homozygous INDELs, which is consistent with the fact that homozygous INDELs are more likely to be detected at a lower coverage shown above (Figure 12B). This shows that one should be cautious about the issue of false-negative heterozygous INDELs in any sequencing experiement with a low coverage (less than 30X). Figure 12B also shows that detection of heterozygous INDELs indeed requires higher coverage than homozygous ones (sensitivity of 95% at 60X vs. 50X). Notably, the number of heterozygous INDELs was 1.6-fold higher than homozygous ones (1600 vs. 635 per sample). This re-affirms the need for 60X mean coverage to achieve a very high accuracy INDEL call set.

## Conclusions

Despite the fact that both WES and WGS have been widely used in biological studies and rare disease diagnosis, limitations of these techniques on INDEL calling are still not well characterized. One reason is that accurate INDEL calling is in general much more difficult than SNP calling. Another reason is that many groups tend to use WES, which we have determined is not ideal for INDEL calling for several reasons. We report here our characterization of calling errors for INDEL detection using Scalpel. As expected, higher coverage improves sensitivity of INDEL calling, and large INDEL detection is uniformly more difficult than detecting smaller INDELs. We also showed that assembly based callers are more capable of revealing a larger spectrum of INDELs, relative to alignment based callers. There are several reasons for the low concordance for WGS and WES on INDEL detection. First, due to the low capture efficiency, WES failed to capture 16% of candidate exons, but even at sites that were successfully captured, there were more coverage biases in the WES data, relative to the WGS data. Second, PCR amplification introduces reads with higher INDEL error rate, especially in regions near homopolymer A/T’s. Lastly, STR regions, especially homopolymer A/T regions were more likely to result in multiple candidates at the same locus. We recommend controlling for homopolymer false INDEL calls with a more stringent filtering criteria. This is essential for population-scale sequencing projects, because the expense of experimental validation scales with the sample size.

Our validation data showed that INDELs called by both WGS and WES data were indeed of high quality and with a low error rate. Even though the WGS data has much lower depth coverage in general, the accuracy of INDEL detection with WGS data is much higher than that with WES data. We also showed that the WES data is missing many large INDELs, which we speculate might be related to the technical challenges of pulling down the molecules containing large INDELs during the exon capture process. Homopolymer A/T INDELs are a major source of low quality INDELs and multiple signature events, and these are highly enriched in the WES data. This was confirmed by the comparison of PCR-free and standard WGS data. In terms of sensitivity, we calculated that WGS at 60X mean coverage from the HiSeq platform is needed to recover 95% of INDELs with Scalpel.

As more and more groups are moving to use new micro-assembly based algorithms such as Scalpel, practical considerations for experimental design should be introduced to the community. Here we present a novel classification scheme utilizing the validation data, and we encourage researchers to use this guideline for evaluating their call sets. The combination of alternative allele coverage and the k-mer Chi-Square score is an effective filter criterion for reducing INDEL calling errors without sacrificing much sensitivity. This classification scheme can be easily applied to screen INDEL calls from all variant callers. Since alternative allele coverage is generally reported in the VCF files, the Chi-Square scores can also be computed directly. For consumer genome sequencing purposes, we recommend sequencing human genomes at a higher coverage with a PCR-free protocol, which can substantially improve the quality of personal genomes. Although this recommendation might initially cost more than the current standard protocol of genome sequencing used by some facilities, we argue that the significantly higher accuracy and decreased costs for validation would ultimately be cost-effective as the sequencing costs continue to decrease, relative to either WES or WGS at a lower coverage. However, it is important to point out that with the release of Illumina HiSeq X-Ten and other newer sequencers, the coverage requirement to accurately detect INDELs may decrease because reads with longer read length can span repetitive regions more easily. Besides, bioinformatics algorithms are another important consideration, and we expect the further enchancements of Scalpel and other algorithms will help reduce the coverage requirement while maintaining a high accuracy.

### List of abbreviations used

(INDELs): Insertions and Deletions
(WGS): whole genome sequencing
(WES): whole exome sequencing
(NGS): next-generation sequencing
bp: (base pair)
PCR: (polymerase chain reaction)
(STR): short tandem repeats
(poly-A): homopolymer A
(poly-C): homopolymer C
(poly-G): homopolymer G
(poly-T): homopolymer T
(STR): short tandem repeats
(other STR): except homopolymers
(poly-A/T): homopolymer A or T

## Authors’ contributions

H.F. analyzed the data and wrote the manuscript. Y.W. optimized the validation experiments and designed the primers. G.N. assisted in characterizing the simulation and validation data. J.A.O. acted as a consultant for the MiSeq validation analyses. Y.W. and L.J.B. performed the Miseq validation experiments. J.R. generated the WGS and WES data. M.R. supervised the generation of the WGS and WES data. I.I. developed the tool for the simulated data. H.F., M.C.S. and G.J.L. designed and analyzed the experiments. G.J.L. developed experimental design for INDEL validation, suggested, reviewed and supervised the data analysis, and wrote the manuscript. All of the authors have read and approved the final manuscript.

## Authors’ information

G.J.L.,M.C.S., M.R. and I.I. are faculty members at Cold Spring Harbor Laboratory (CSHL). G.N was a post-doctoral fellow at CSHL and is currently employed at the New York Genome Center. J.R. is a laboratory technician at CSHL. H.F., J.A.O., and Y.W. are graduate students at CSHL and Stony Brook University. L.J.B. is a visiting undergraduate student at CSHL and a undergraduate student at Universidad Nacional Autonoma de Mexico.

## Acknowledgements

The laboratory of G.J.L. is supported by funds from the Stanley Institute for Cognitive Genomics at Cold Spring Harbor Laboratory (CSHL). The laboratory of M.C.S. is supported, in part, by National Institutes of Health award (R01-HG006677) and by National Science Foundation award (DBI-1350041). The CSHL genome center is supported in part by a Cancer Center Support Grant (CA045508) from the NCI. This work was partially supported by a grant from the Simons Foundation (SF235988) to Michael Wigler. We are grateful to all of the families at the participating SFARI Simplex Collection (SSC) sites, as well as the principal investigators (A. Beaudet, R. Bernier, J. Constantino, E. Cook, E. Fombonne, D. Geschwind, D. Grice, A. Klin, R. Kochel, D. Ledbetter, C. Lord, C. Martin, D. Martin, R. Maxim, J. Miles, O. Ousley, B. Pelphrey, B. Peterson, J. Piggot, C. Saulnier, M. State, W. Stone, J. Sutcliffe, C. Walsh, and E. Wijsman). The DNA samples used in this work are included within SSC release 15. Approved researchers can obtain the SSC population dataset described in this study by applying at https://base.sfari.org. We thank Sara Ballouz, Wim Verleyen, Jesse Gillis, Ruibang Luo, Shane McCarthy, Zamin Iqbal, David Mittelman, Martin Reese, and the anonymous reviewers for helpful discussions and comments on the paper.

## Additional files

### Additional file 1 – Supplemental figures and tables

This file includes supplemental figure S1-S4, supplemental table S1-S9, and their corresponding figure/table legends. This file also includes supplemental note 1 describing the analysis of *de novo* INDEL calls.

